# Evolutionary dynamics at the leading edge of biological invasions

**DOI:** 10.1101/2024.12.13.628423

**Authors:** Silas Poloni, Frithjof Lutscher, Mark A. Lewis

## Abstract

Empirical evidence shows that evolution may take place during species’ range expansion. Indeed, dispersal ability tends to be selected for at the leading edge of invasions, ultimately increasing a species’ spreading speed. However, for organisms across many different taxa, higher dispersal comes at the cost of fitness, producing evolutionary trade-offs at the leading edge. Using reaction-diffusion equations and adaptive dynamics, we provide new insights on how such evolutionary processes take place. We show how evolution may drive phenotypes at the leading edge to maximize the asymptotic spreading speed, and conditions under which phenotypic plasticity in dispersal is selected for under different dispersal-reproduction trade-off scenarios. We provide some possible future research directions and other systems where the framework can be applied.

## 1 Introduction

Novel empirical evidence shows that rapid adaptation of dispersal and demographic traits can take place during range expansion processes [Shine et al., 2011, Ochocki and Miller, 2017, Weiss-Lehman et al., 2017]. The evolutionary and ecological consequences of such adaptations are many, and they impose novel challenges in our ability to measure and forecast the future range of a given invasive species, on both experimental and theoretical grounds [Miller et al., 2020, Lustenhouwer et al., 2023]. In this work, we develop new theory for outcomes arising from evolutionary trade-offs at the leading edge of an invasion. Recent work on mathematical models of trait-structured populations can explain how individual variability in dispersal and growth can change the speed of invasion [Elliott and Cornell, 2012, Keenan and Cornell, 2021] and trait distributions at the leading edge [Poloni and Lutscher, 2023], but the connection between those results has not been fully discussed or mathematically formalized. In this work, we present a new framework of analysis for understanding how speeds of invasion and leading-edge trait distributions are connected.

During range expansion, space itself becomes a driver of evolution and selection. Individuals that are able to colonize novel regions faster tend to experience less competitive pressure from con-specifics when they spread into low-density regions. These individuals breed, on average, individuals equally capable of quickly spreading, and the best spreaders continue to be selected. The process repeats and gives rise to what is now referred to as spatial sorting, with phenotypes being organized in space from least spreading abilities at the core of invasion to best spreading abilities at the leading edge [Shine et al., 2011, Phillips and Perkins, 2019]. This highlights the existence of different evolutionary forces at play throughout space, since phenotypes that favor spread are expected to be selected for at the leading edge, whereas at the core of invasion, selection for phenotypes that help reduce competitive pressure is expected to take place [Burton et al., 2010, Perkins et al., 2013] (but see Shaw et al. [2023]).

Nonetheless, phenotypes related to dispersal and reproduction, both key for the spread of a species, usually present trade-off relations. For example, different insect taxa display negative correlations between fecundity and flight capacity [Hanski et al., 2006, Elliott and Evenden, 2012, Tigreros and Davidowitz, 2019]. Plants that reproduce and disperse through wind-borne seeds may present trade-offs between nutrient load and distance traveled per seed [Greene and Johnson, 1993, Eriksson and Jakobsson, 1999]. Cane toads that display larger and strong back legs also present lower gonad mass Kelehear and Shine [2020]. Under such trade-offs, individuals at the leading edge are potentially under different evolutionary forces, pushing for higher dispersal and pulling for higher reproductive ability Phillips and Perkins [2019].

Such evolutionary processes challenge us to understand biological invasions via experiments Sherpa and Després [2021]. Experiments usually require long spatial and temporal scales, and may present a high risk of unintended spread of studied organisms in non-native habitats, leaving us with studies from successful past invasions, in which some evolutionary aspects may have already eloped. However, novel experiments in laboratory environments (on relatively small spatial scales) show that such evolutionary processes play an important role on speeding up invasions, due to adaptations leading to higher dispersal and/or growth rates of the invading species [Ochocki and Miller, 2017, Weiss-Lehman et al., 2017, Szűcs et al., 2017, Deforet et al., 2019].

Alongside novel experiments, theoretical approaches have been developed to understand how individual variability in dispersal and growth can lead to different outcomes in range expansion. Elliott and Cornell [2012] show that dimorphic populations can, under some conditions, travel faster than any of their composing morphs would be able to individually. They refer to this phenomenon as a population presenting an *anomalous spreading speed*. The appearance of such speeds requires a significant growth and dispersal trade-off between the two different morphs and some level (albeit really small) of imperfect trait inheritance among them. Later, Keenan and Cornell [2021] extended the theory for any given finite number of morphs and provided the exact reproduction and dispersal trade-off conditions under which anomalous spreading speeds appear.

Through a different modeling approach, Poloni and Lutscher [2023] showed that when anomalous spreading speeds appear (under similar conditions as in Keenan and Cornell [2021]), population trait distributions at the leading edge tend to be bimodal, with one mode at maximum dispersal and minimum growth, and the other at the opposite end of the trait spectrum, representing a dimorphism at the leading edge. Whenever anomalous spreading speeds did not appear, trait distributions at the leading edge tended to be unimodal, with a single-trait speed maximized. These results help connect population spreading speeds to leading edge trait distributions, and together, reveal the evolutionary forces at play: selection at the leading edge giving rise to phenotypes that maximize spreading speeds, in a similar fashion as found in Phillips and Perkins [2019].

The connection between such results and concepts, however, remains to be formalized. Here, we aim to do so using the adaptive dynamics framework Geritz et al. [1998], Diekmann [2003]. In adaptive dynamics, a resident population is assumed to be at ecological equilibrium when a mutant with a novel phenotype appears. The mutant’s fate is determined by its growth rate in an environment set by the resident population. This information is usually depicted in a pairwise invasibility plot, a diagram indicating whether the mutant’s phenotype provide a fitness gain compared to resident’s phenotype. Deforet et al. [2019] established criterion for successful mutant invasion at the leading edge, and contrast it with experimental results for two strains of the bacteria *P. aeruginosa*. Here, using linearization analysis, we find a similar criterion as found in Deforet et al. [2019], and with it, apply the adaptive dynamics framework to provide novel ways to understand such evolutionary processes.

The paper is organized as follows: In section 2, we present the modeling approach, assumptions taken and formal method of analysis. We find the same criteria as Deforet et al. [2019] and explore examples in section 3. We not only recover results from Keenan and Cornell [2021], Poloni and Lutscher [2023], but also provide new insights on how such results come to be. Finally, in section 4, we discuss our results and provide possible avenues of future research.

## 2 Model and analysis framework

We use the reaction-diffusion modeling framework to track different morphs of an invading population in space and time [Cantrell and Cosner, 2004, Lewis et al., 2016]. We denote the density of the *i*-th morph in space (*x*) and at time (*t*) as *u*_*i* ≡_ *u*_*i*_(*x, t*). All morphs compete for a shared resource, such that we can measure their densities in terms of the species’ carrying capacity. Each morph is characterized by its diffusion coefficient, *D*_*i*_, and its growth rate, *r*_*i*_.

To account for trade-offs between dispersal and reproduction, we assume that growth and dispersal are negatively correlated, such that the intrinsic growth rate of a morph depends on its diffusion coefficient, i.e., *r*_*i* ≡_ *r*(*D*_*i*_) > 0, with *r*′(*D*_*i*_) < 0, i.e., *r* is decreasing with *D*_*i*_. Trade-offs can display concave-up or down shapes, depending precisely on how the reproduction and dispersal phenotypes are anti-correlated. In concave-up (resp. concave-down) trade-offs, increases from small dispersal do (resp. do not) lead to a significant loss in reproduction.

First, to simplify analysis and enhance clarity, let the invading species be monomorphic upon introduction, with diffusion coefficient *D*_1_ and growth rate *r*_1_ = *r*(*D*_1_). This leads to the classical Fisher-Kolmogorov equation [Fisher, 1937, Kolmogorov et al., 1937]:

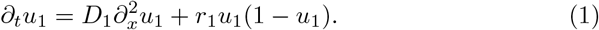

Equation (1) admits traveling wave solutions and presents a linearly determinate asymptotic spreading speed, i.e., the speed of spread of the population depends on leading edge dynamics, where the population is at low density ^1^ [Weinberger, 1982].

The classical expression for the asymptotic spreading speed for Equation (1) is 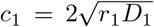. For compact initial data, the speed *c*_1_ is only achieved at the asymptotic limit *t* → *∞*. However, the limit is approached with reasonable precision after a relatively short transient phase Kolmogorov et al. [1937], Kanel’ [1962], Fife and McLeod [1977]. It will be useful to define

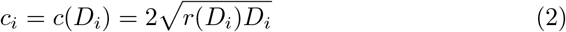

as the asymptotic spreading speed of a morph with diffusion coefficient *D*_*i*_.

Now, assume that a mutant morph is born at the leading edge, after the transient phase between early invasion and convergence towards a traveling wave formation has passed. The mutant has diffusion coefficient *D*_2_, and, consequently, growth rate *r*_2_ = *r*(*D*_2_). We will be concerned about whether this mutant morph is able to stay at the leading edge and replace the resident morph therein. If so, will the process of repeated introductions of mutant morphs ever come to a halt? If so, will this lead to the establishment of the fastest possible morph at the expansion front of the species?

To answer this question, we will deploy ideas from the framework of adaptive dynamics [Geritz et al., 1998, Diekmann, 2003], which, similar to our goal here, addresses how a given trait in a population evolves through consecutive favorable mutations. However, adaptive dynamics requires the resident population to be at ecological equilibrium, assuming a time scale separation between ecological and evolutionary phenomena. Here, although the population is not at ecological equilibrium, the front of the leading edge remains unchanged from the perspective of an observer who travels alongside it. For that reason, a traveling wave is sometimes referred to as a “relative equilibrium”.

Importantly, we consider two different types of mutant populations. First, we analyze the case where mutants possess novel phenotypes, different than those of the resident population, and both resident and mutant reproduction is perfectly clonal, i.e., parent and offspring have the exact same phenotypes. This is the classical notion of adaptive dynamics Geritz et al. [1998]. Second, we study the case where mutants can exhibit both novel and resident phenotypes, i.e., mutation is not on the phenotype itself, but on the ability to express said phenotypes, yielding a mutant population that possesses some phenotypic plasticity. The mutant population therefore can exhibit imperfect clonal reproduction, with their offspring having either resident or novel phenotypes. This last case might be particularly important for species that can display polymorphisms mediated by different phenotypic inheritance mechanisms, for example epigenetics and polyploidy, leading to what is more broadly referred to as phenotypic plasticity Bonduriansky and Day [2020], Adams and Wendel [2005].

### 2.1 Perfect clonal reproduction

Consider the case where offspring phenotypes are identical to those of their parents. Let *u*_1_ and *u*_2_ be the densities of the resident and mutant populations, respectively. In this case, the governing equations for both morphs are the diffusive competitive Lotka-Voltera equations:

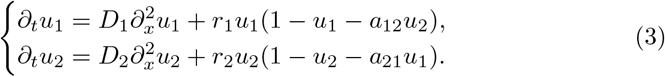

However, since we assume that the mutant appears at the leading edge of a traveling wave composed of the resident population, both *u*_1_ and *u*_2_ are small, since mutants are rare. The equations decouple, and we can analyze them as the release of two non-interacting morphs. The new leading edge is therefore composed of the fastest morph, i.e., the one with the highest asymptotic spreading speed. We represent the process graphically in figure 1, using a numerical simulation, and provide details in the figure legend.

**Figure 1.**
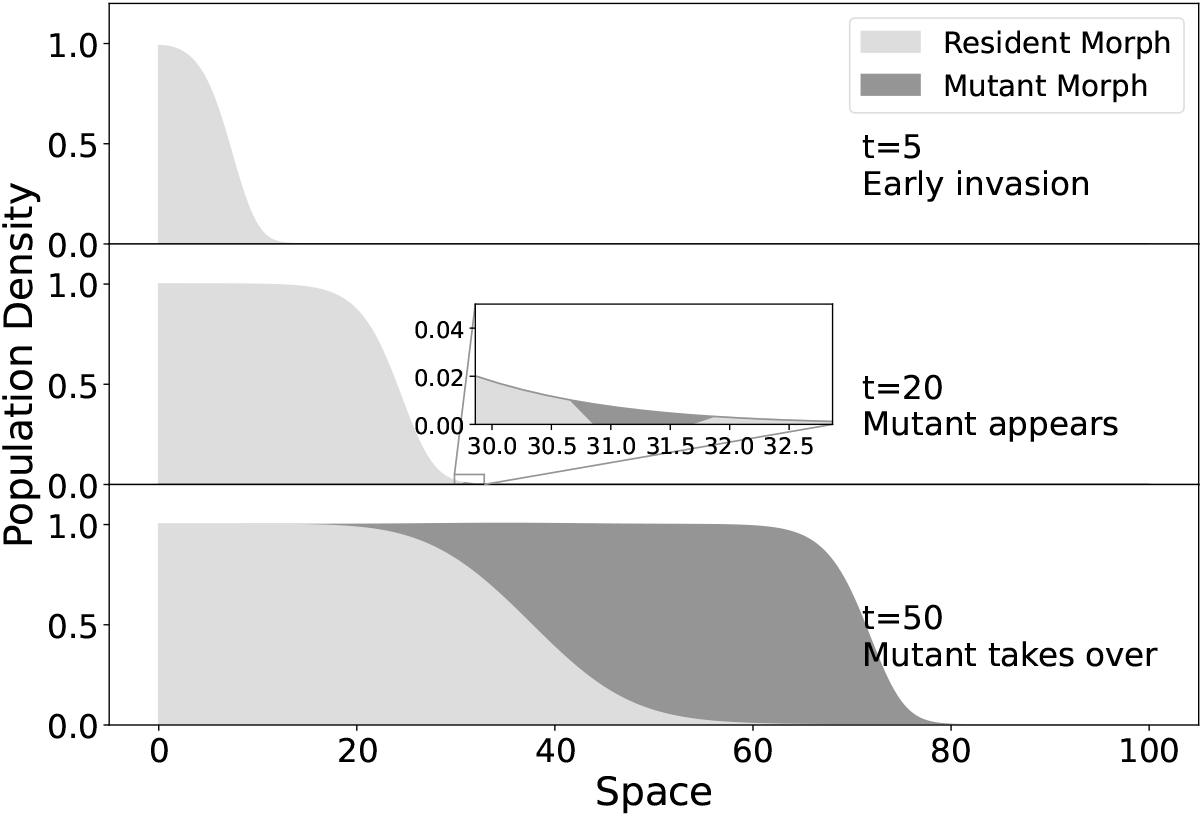
An illustration of a successful mutant in the perfect clonal reproduction case. A monomorphic resident population starts spreading (top). After some time elapses, the resident population attains a traveling wave profile with speed *c*(*D*_1_), and a small mutant density appears at the leading edge (middle). Later, the mutant population takes over the leading edge and spreads at speed *c*(*D*_2_) *> c*(*D*_1_), forcing the observer to adjust its speed to keep up with the invading population (bottom).

This way, morph 2 invades if *c*(*D*_2_) *> c*(*D*_1_). Note that replacement occurs only at the leading edge, and is independent of how competition between these two morphs plays out at the core and in the wake of the invasion. In fact, if the resident morph is the better competitor (*a*_12_ < 1 *< a*_21_), it will spread into the mutant population’s territory and replace them at speed 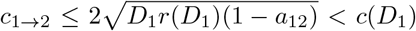, however, since *c*(*D*_1_) *< c*(*D*_2_), the resident will never catch up with mutant population while the range expansion process lasts [Hosono, 1998, Lewis et al., 2002, Weinberger et al., 2002].

To put this result into the classical perspective of adaptive dynamics, we denote by *s*(*D*_1_, *D*_2_) the invasion exponent, defined as the expected growth rate of the mutant population with phenotype *D*_2_ in an environment set by a resident population with phenotype *D*_1_ Diekmann [2003]. This way, *s*(*D*_1_, *D*_2_) > 0 implies successful mutant invasion, while *s*(*D*_1_, *D*_2_) < 0 implies its failure. The precise mathematical definition of *s*(*D*_1_, *D*_2_) is given by an eigenvalue problem, and its precise functional form is found by solving said eigenvalue problem (supplemental material). We don’t require the latter to proceed, and infer that *s*(*D*_1_, *D*_2_) is positive when *c*(*D*_2_) *> c*(*D*_1_), and negative otherwise, as aforementioned.

Hence, we can write

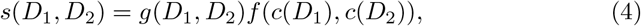

with *g* = *g*(*D*_1_, *D*_2_) > 0 and *f* (*c, c*) = 0 for all *c >* 0. Here, we assume that *f* = *f* (*c*_1_, *c*_2_) is smooth and twice differentiable near *c*_1_ = *c*_2_ (and *D*_1_ = *D*_2_ by consequence). Furthermore, *f* is positive if *c*_2_ *> c*_1_ and negative if *c*_2_ *< c*_1_, and its is decreasing (resp. increasing) in the first (resp. second) argument near the line *c*_1_ = *c*_2_ (by construction). The successful invasion criterion is sign(*s*(*D*_1_, *D*_2_)) = sign(*f*) > 0, so morphs that have higher speeds invade morphs with lower speeds at the leading edge.

In adaptive dynamics, we assume that mutations lead to small changes in traits, such that the invasion exponent can be written as

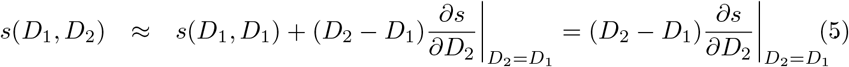

Therefore, amutant population grows if the first derivative of *s* with respect to *D*_2_ is positive, and declines otherwise. The first derivative of the invasion exponent w.r.t. a mutant’s phenotype, 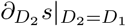, is also called the selection gradient, as it gives the direction of increase in mutant growth rate, and hence the direction evolution will push phenotypes to.

The selection gradient with *s* as defined in (4) is

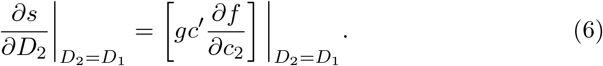

Since *f* is increasing with *c*_2_ near *D*_1_ = *D*_2_, the sign of the selection gradient is the same as the sign of *c*′(*D*_1_), i.e., the direction of evolution follows the increase of the species’ asymptotic spreading speed. Therefore, a singular strategy, *D*^∗^, for which we have 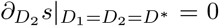, is also a critical point of *c*. Using the expression 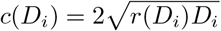, we can find D∗ by solving

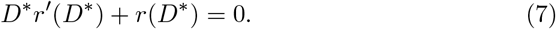

Note that equation (7) cannot be satisfied if *r* is monotonically increasing with *D*_*i*_, i.e., without trade-offs. In that case, evolution would keep pushing morphs at the leading edge towards higher dispersal ability and growth rates.

The singular strategy *D*^∗^ is called an evolutionary stable strategy (ESS) if all mutants fail to invade a resident population with strategy *D*^∗^, i.e., if *D*^∗^ is a maximum of *s*(*D*^∗^, *D*_2_). *D*^∗^ is an ESS if

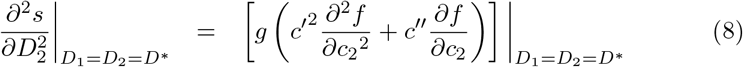

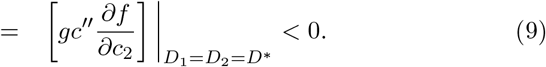

Again, since *f* is increasing, this condition is only satisfied if *c*^′′^(*D*^∗^) < 0. As expected, an ESS is such that it maximizes the species’ asymptotic spreading speed.

When the singular strategy, *D*^∗^, is not an ESS, we have *c*^′′^ (*D*^∗^) > 0, i.e., *D*^∗^ is a minimum of *c*. Because 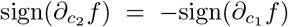 it follows that 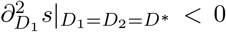, and evolution will drive phenotypes away from *D*^∗^. In this case, the strategy *D*^∗^ is referred to as an evolutionary repeller (see figure 14 in Diekmann [2003]).

If *D*^∗^ is an ESS, it is important to address whether it can be gradually approached by subsequent mutant invasions closer to *D*^∗^, i.e., whether *D*^∗^ is a convergent stable strategy. For that, we must have that strategies *D*_2_ ∈ (*D*_1_, *D*^∗^) (resp. *D*_2_ ∈ (*D*^∗^, *D*_1_)) successfully invade strategies *D*_1_ *< D*^∗^ (resp. *D*_1_ *> D*^∗^). Therefore, in the vicinity of *D*_1_ = *D*_2_ = *D*^∗^, the selection gradient must be positive in *D*_1_ *< D*^∗^ and negative in *D*_1_ *> D*^∗^, or more generally,

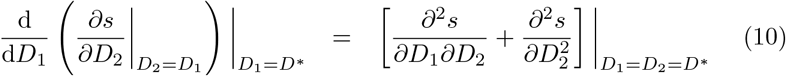

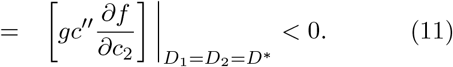

Therefore, the condition for *D*^∗^ to be an evolutionary and a convergent stable strategy are the same. When a strategy is both evolutionarily and convergent stable, it is classically addressed as a continuous stable strategy (CSS) [Diekmann, 2003]. In terms of the growth function *r*(*D*_*i*_), the singular strategy *D*^∗^ is a CSS if

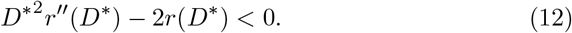

Note that if a singular strategy *D*^∗^ exists and *r*(*D*_*i*_) is concave down, i.e., *r*^′′^ < 0, equation (12) is always satisfied and *D*^∗^ is a CSS. Therefore, concave up (convex) trade-offs are necessary, but not sufficient, to produce evolutionary repellers.

We still need to check whether dimorphisms are possible near the singular strategy *D*^∗^. For that, we need a pair of strategies *D*_1_, *D*_2_, with *D*_1_ *< D*^∗^ *< D*_2_, such that *s*(*D*_1_, *D*_2_) > 0 and *s*(*D*_2_, *D*_1_) > 0, so that they are mutually invasible. This is impossible, since sign(*s*(*D*_2_, *D*_1_)) = −sign(*s*(*D*_1_, *D*_2_)). It is important to address the exceptional case where two morphs with precisely the same spreading speed evolve away from a repeller strategy *D*^∗^. In this scenario, both morphs can coexist at the leading edge, however, any further adaptation from either morph will disrupt the dimorphism, leading again to a monomorphic state.

### 2.2 Imperfect clonal reproduction

Now, we turn our attention to the case where mutants can express both the resident and a novel phenotype. This leads to the following system of equations

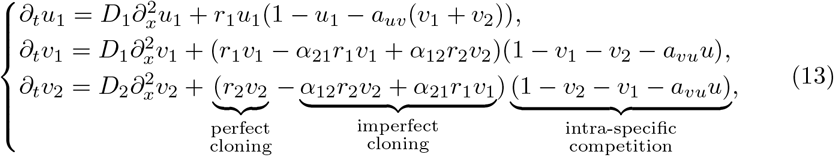

where *u*_1_ is the resident population and *v*_1_ and *v*_2_ are mutant populations with phenotypes *D*_1_ and *D*_2_, respectively. Competition between resident and different mutants is expressed in terms of the coefficients *a*_*uv*_ and *a*_*vu*_. The *α*_*ij*_ ∈ (0, 1), *i, j* = 1, 2, *j* ≠ *i*, are the proportion of mutant offspring from parents of phenotype *j* born with phenotype *i*.

Linearizing the model around the leading edge, we find that, similarly to the perfect reproduction case, mutants invade the leading edge if they have a higher asymptotic spreading speed. However, the conditions for that are slightly more intricate than before.

Following Elliott and Cornell [2012], we define the set

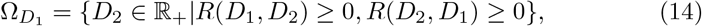

where

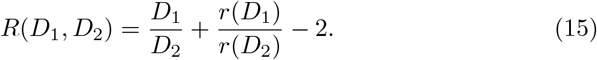

If a mutant morph expresses a phenotype 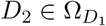 (and therefore 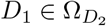), then mutants can spread with the *anomalous spreading speed*

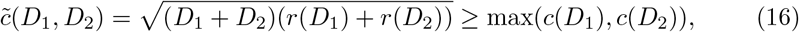

at least in the limit *α*_*ij*_ ≪ 1, which is the main result from Elliott and Cornell [2012], Keenan and Cornell [2021]. This, in turn, implies that the mutant invades the leading edge and strongly displays both *D*_1_ and *D*_2_, leading to dimorphism [Poloni and Lutscher, 2023]. The process is depicted in figure 2, with a numerical simulation (details in figure caption).

**Figure 2.**
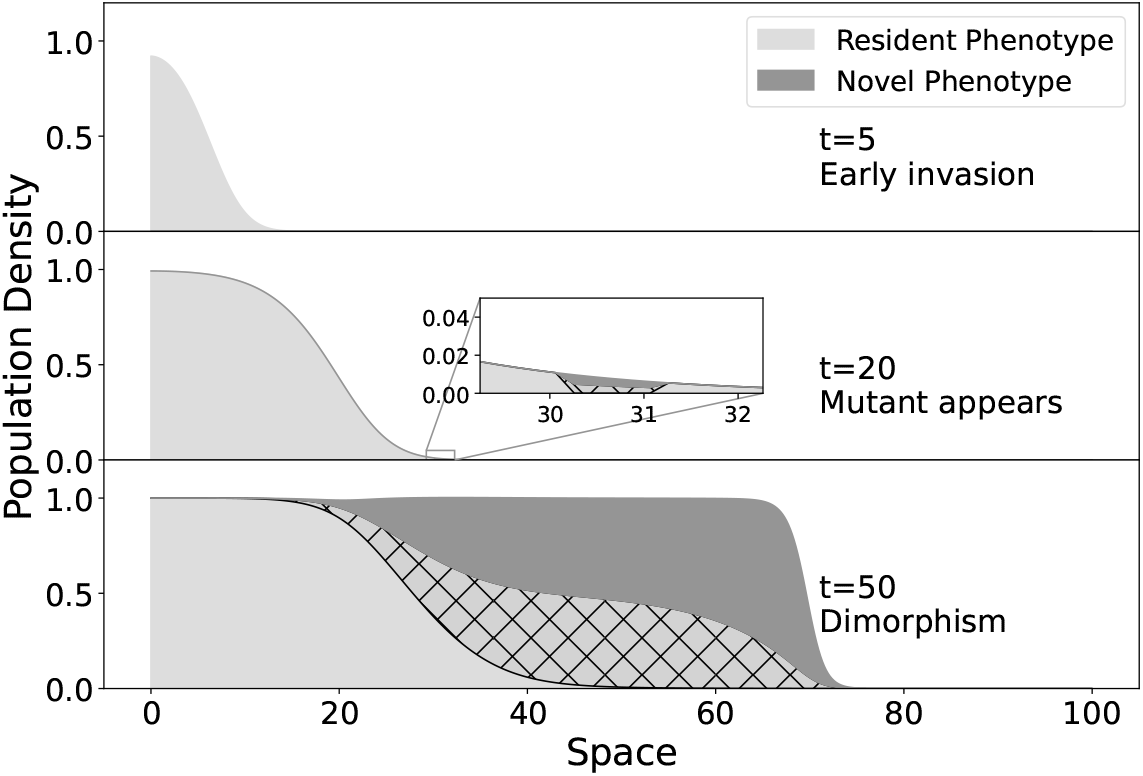
An illustration of a successful mutant in the imperfect clonal reproduction case. A monomorphic resident population starts spreading (top). After some time elapses, at the middle panel, the resident population attains a traveling wave profile with speed *c*(*D*_1_), and a small mutant density appears at the leading edge, displaying both resident and novel phenotypes (middle). Later, both novel and resident phenotypes are present at the leading edge, spreading at speed 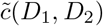 (bottom). The dashed region represents the portion of mutant population expressing resident phenotypes.

Note that, although the densities of both mutant morphs at the leading edge are positive (since *α*_*ij*_ > 0), their densities are of comparable order only if 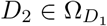. When the mutant expresses a phenotype 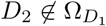, the problem is similar to the case of perfect clonal reproduction in that only the fastest phenotype is present at the leading edge. Note that in this case, selection is not only for the fastest phenotype but also against the expression of the slowest phenotype altogether. This way, if *c*(*D*_2_) *> c*(*D*_1_), mutants can establish, but will predominantly display *D*_2_ at the leading edge, otherwise, mutants fail to invade. Expressing *D*_2_ reduces the spreading speed of mutants, so that the expression mechanism is lost and the resident population stays at the leading edge alone. We provide an extensive mathematical description of these cases in the supplemental material.

Therefore, we focus on mutants with phenotypes 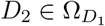, that successfully invade the leading edge forming a dimorphic population. A valid question is where will evolution take the phenotypes of such a dimorphic population once it is established? To study this problem, we let a resident pair of morphs with strategies *D*_1_, *D*_2_ be invaded by a pair of mutants with strategies *d*_1_, *d*_2_. We assume that a mutant pair invades the resident pair if 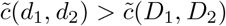. As before, we define

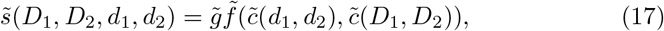

where 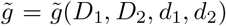 and 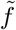 have the same properties as *g* and *f* in (4), respectively.

Since the speed of the mutant morphs does not depend on the resident phenotypes, a singular strategy is a critical point 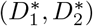 of 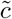, that can be found by solving

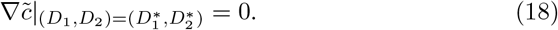

Such singular strategies can then be classified by analyzing the Hessian matrix of 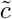. The existence of solutions 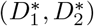 to this equation depend on the specific reproduction-dispersal trade-off, *r*(*D*_*i*_).

The approach here is slightly different from what was presented so far, in the sense that we allow mutations to take place in both morphs simultaneously. Clearly, we do not expect simultaneous mutations, since they are rare and could occur in each of the morphs independently. However, this abstraction helps us visualize the direction in which phenotypes are expected to evolve at the leading edge, as we will see in section 3.

We start our analysis by noticing that on the line *D*_1_ = *D*_2_, we have 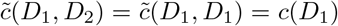. A quick calculus exercise shows that a critical point *D*^∗^ of *c*, is also a critical point *D*_1_ = *D*_2_ = *D*^∗^ of 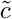. We must therefore determine whether *D*^∗^ is an attractor or repeller of both monomorphic and dimorphic populations or a saddle point, meaning it can be approached monomorphically but not dimorphically or vice-versa.

#### Continuously stable strategy

We call (*D*_1_, *D*_2_) = (*D*^∗^, *D*^∗^) a CSS (both an evolutionary and a convergent stable strategy) if it is a local maximum of both the anomalous asymptotic spreading speed and the monomorphic asymptotic spreading speed, i.e., if

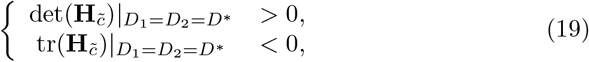

where 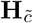 is the Hessian matrix of 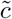. A quick calculation shows that

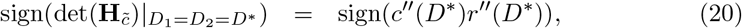

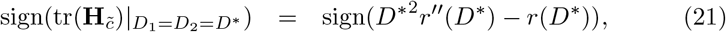

which alongside equation (12) simplifies condition (19) to

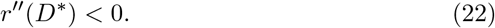

Therefore, if a singular strategy exists and the trade-off is concave-down, then the singular strategy is a CSS. This implies that during the course of invasion, both initially monomorphic and dimorphic populations with phenotypes close to *D*^∗^ will evolve towards *D*^∗^ at the leading edge.

#### Evolutionary repeller

The singular strategy (*D*_1_, *D*_2_) = (*D*^∗^, *D*^∗^) is an evolutionary repeller if it is a local minimum of both monomorphic (*c*) and dimorphic 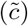 spreading speeds. That is, if

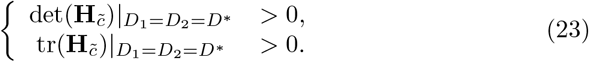

The conditions on the trace of 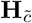 can be omitted in view of (12) and (21). Alongside (20), the condition on the determinant simplifies, since *r*^′′^(*D*^∗^) < 0 ⟹ *c*^′′^(*D*^∗^) < 0, leading us to the condition

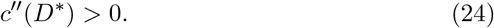

Noticeably, the condition for a singular strategy to be an evolutionary repeller is the same in the perfect and imperfect clonal reproduction cases. Here, initially monomorphic (dimorphic) populations would remain monomorphic (dimorphic) and evolve away from *D*^∗^.

#### Evolutionary saddle points

In the case where *D*^∗^ is a saddle point, we must have

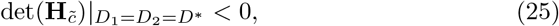

with *D*^∗^ being a local maximum of *c*(*D*_*i*_), since when it is the minimum, *D*^∗^ is an evolutionary repeller. We therefore have the simplified conditions

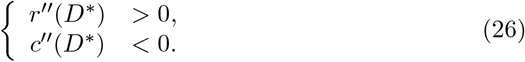

A saddle point can only be an attractor of initially monomorphic invasions. Once the species is at strategy *D*^∗^, any further phenotypic variation that appears at the leading edge is maintained, forming a dimorphic pair with strategies that both evolve away from *D*^∗^ as invasion progresses.

While it is tempting to refer to this saddle point as a branching point, we should avoid the misconception. Both morphs can only remain at the leading edge if there is a mechanism that allows for a relatively fast switch in dispersal phenotypes. This way, to avoid the misconception, we will refer to such a point as an evolutionary saddle point. In such points, what is being selected for is the maintenance of the mechanism for dimorphisms, rather than the morphs themselves.

Such points are unlikely to appear in the classical analysis of adaptive dynamics, because neither of the morphs would be able to invade alone. That is because, when the resident is established with an optimal phenotype *D*^∗^, we have *s*(*D*^∗^, *d*_*i*_) = *gf* (*c*(*d*_*i*_), *c*(*D*^∗^)) < 0 ∀*d*_*i*_ ≠ *D*^∗^. However, for a mutant population displaying strategies *d*_1_ and *d*_2_, such that it spreads at an anomalous spreading speed, mutant invasion is possible. In this case, we evaluate 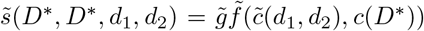, which is positive when *D*^∗^ is a saddle point. This way, such points are possibly only present in spatial contexts where anomalous spreading speeds can emerge, or when advantages of phenotypic plasticity are explicitly accounted for (we provide some mathematical analysis of such cases in the supplemental material).

Importantly, in evolutionary saddle points, because the local maximum is achieved over the line *D*_1_ = *D*_2_, the local minimum is achieved over the perpendicular line that passes through *D*^∗^, *D*_2_ = 2*D*^∗^ − *D*_1_. The direction of maximum increase in dimorphic spreading speeds is also along that line. Therefore, extremes of high (resp. low) dispersal and low (resp. high) growth rates are continuously selected for at the leading edge. Clearly, such strategies are unrealistic and other sources of selection, such as phenotype extremes being associated with high mortality, would prevent these extremes from being attained. In our analysis, these would imply different critical points of 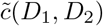, beyond *D*_1_ = *D*_2_ = *D*^∗^. We won’t cover those here, but see Keenan and Cornell [2021] figure 3f and Poloni and Lutscher [2023] figure 8, for the matching case in terms of spreading speeds and leading edge distributions, respectively.

**Figure 3.**
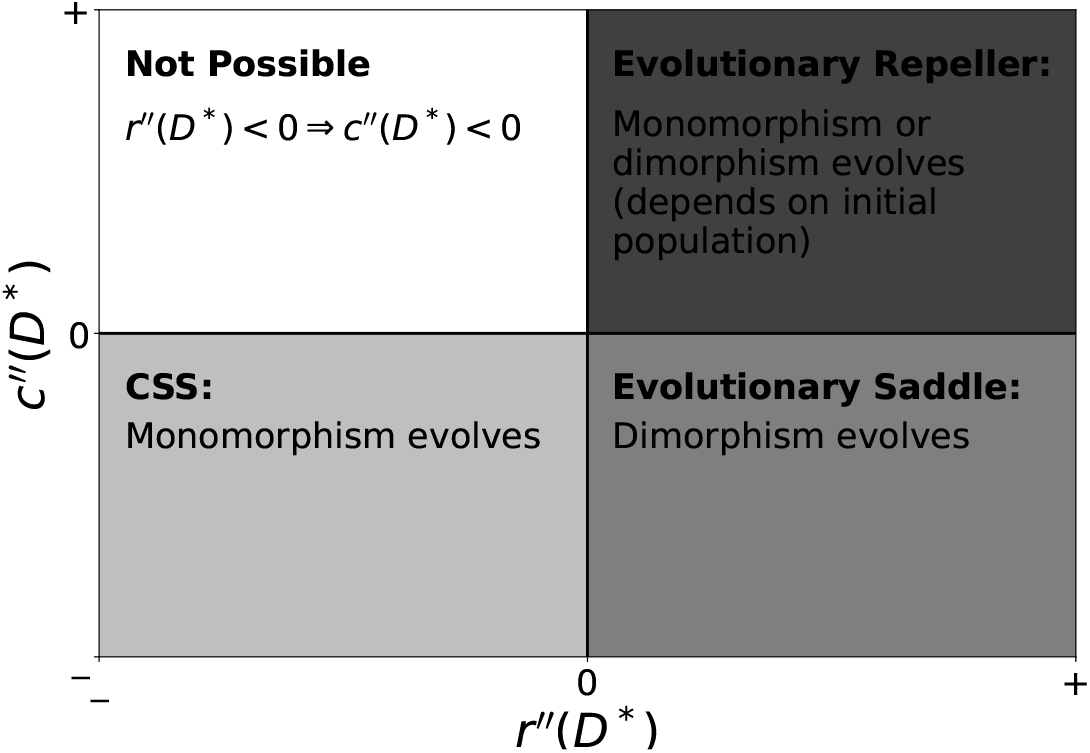
Summary of possible evolutionary outcomes given a singular strategy *D*_1_ = *D*_2_ = *D*^∗^ based on the signs of *r*^′′^(*D*^∗^) and *c*^′′^(*D*^∗^). The white region is not a possible configuration because *r*^′′^(*D*^∗^) < 0 ⇒ *c*^′′^(*D*^∗^) < 0.

#### 2.2.1 Summary of evolutionary outcomes

As we just showed, concavity of the reproduction-dispersal trade-off curve plays an important role on the qualitative behavior of a singular strategy. A concavedown trade-off will lead to a CSS, and selection for a monomorphic population at the leading edge occurs, whereas a concave-up trade off can lead to dimorphism, either via an evolutionary saddle point (for a concave-down *c*(*D*_*i*_)) or an evolutionary repeller (concave-up *c*(*D*_*i*_)). All different evolutionary outcomes of a singular strategy *D*_1_ = *D*_2_ = *D*^∗^ are also summarized in figure 3.

In concave-down trade-offs, intermediate dispersal abilities do not impact reproductive outcomes heavily. Therefore, an optimal combination of *r*(*D*_*i*_) and *D*_*i*_ is selected for at the leading edge. In concave-up trade-offs, on the other hand, such combinations are impossible, since intermediate dispersal abilities cause a significant loss in individual reproductive outcome. Hence, selection at the leading edge pushes the species to the extremes of high dispersal/low reproduction and low dispersal/high reproduction, either forming monomorphic or dimorphic populations.

## 3 Examples

In this section, we will present some reproduction and dispersal trade-off expressions, and the pairwise invasibility plots to which they lead. These different trade-offs can also illustrate how dispersal and reproductive phenotypes of several organisms may evolve during range expansion. Here, we only illustrate possible evolutionary outcomes through modeling examples, without focusing on a specific set of organisms. In all plots we set *g* = 1 and *f* = *c*_2_ − *c*_1_ (and similarly to 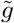 and 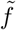).

### 3.1 Continuously Stable Strategies and Evolutionary Saddle points

Following Marculis et al. [2020], we consider a trade-off between growth/reproduction and dispersal modeled by the following expression

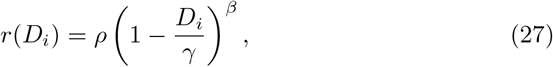

where *D*_*i*_ ∈ (0, *γ*), *γ* is a limiting diffusion coefficient (after which growth rates become negative), ρ > 0 gives the proper dimensions for the growth rate and *β >* 0 is a shape parameter, setting *r* to be concave-down in case *β <* 1 and concave-up if *β >* 1.

The asymptotic spreading speed is found by substituting equation (27) into (2), and the singular strategy is

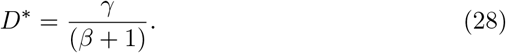

Note that *D*^∗^ is an intermediate value (0 *< D*^∗^ *< γ*) that maximizes *D*_*i*_*r*(*D*_*i*_). The singular strategy maximizes monormphic spreading speeds by achieving intermediate diffusion coefficients and growth rates.

In the case of perfect clonal reproduction, the singular strategy *D*^∗^ is a CSS, since

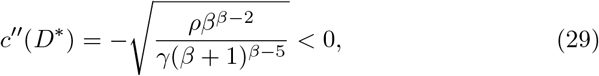

which is also represented in the pairwise invasibility plot in figure 4. Note that resident strategies *D*_1_ *< D*^∗^ are invaded by mutant strategies *D*_2_ ∈ (*D*_1_, *D*^∗^], while resident strategies *D*_1_ *> D*^∗^ are invaded by mutant strategies *D*_2_ ∈ [*D*^∗^, *D*_1_). These are both represented by the dark gray areas, also signaled with a plus sign (+). The vertical dashed line that passes through *D*^∗^ lies entirely on the light gray region, signaling that once *D*^∗^ is established, it cannot be invaded by any other mutant *D*_2_. Likewise, the horizontal dashed line lies entirely on the dark gray region, signaling that mutants with strategy *D*_2_ = *D*^∗^ invade any other resident strategy *D*_1_. This implies, subsequent successful mutations will lead phenotypes at the leading edge closer to *D*^∗^, increasing the species invasion speed until *D*^∗^ is attained.

**Figure 4.**
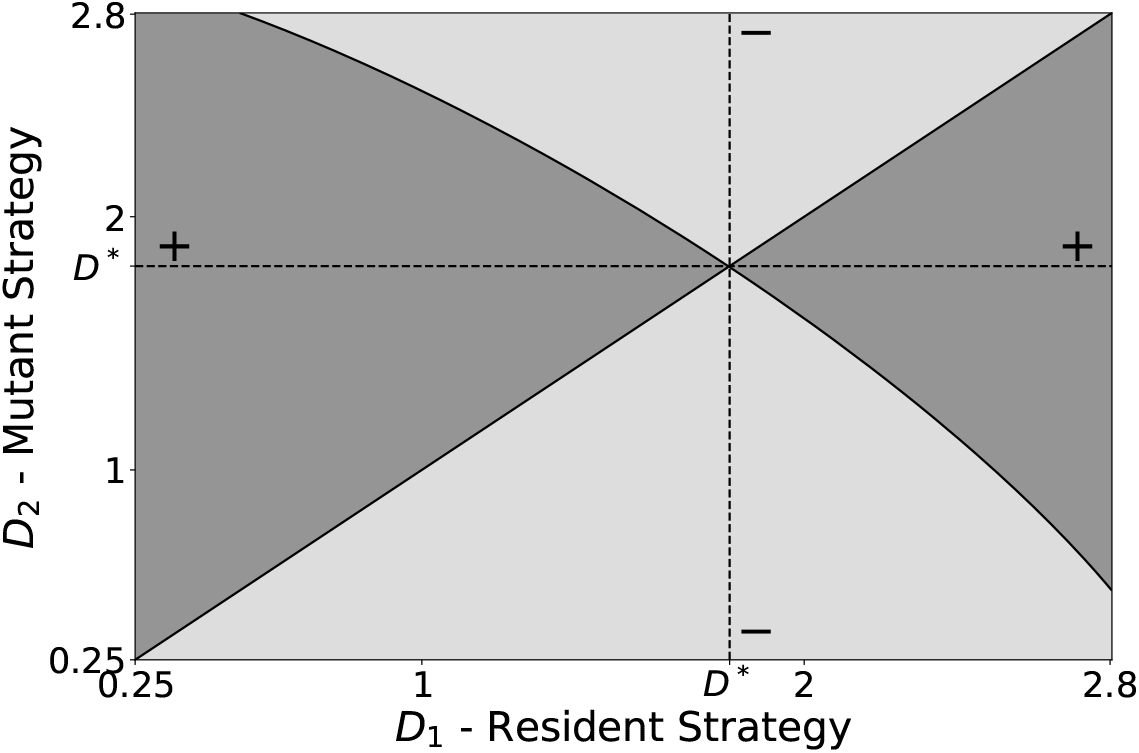
The pairwise invasibility plot indicates that *D*^∗^ = *γ/*(*β* + 1) is a CSS in the perfect clonal reproduction case, with *r*(*D*_*i*_) as defined by equation (27). Light and dark gray regions show where mutants *D*_2_ fail and succeed to invade residents *D*_1_, respectively. Parameters used are *ρ* = *γ* = 3 and *β* = 2*/*3.

From the perfect clonal reproduction analysis, we have that *D*^∗^ is a CSS, meaning it is approached monomorphically. This way, in the imperfect clonal reproduction case, where dimorphisms are possible, we can only have that *D*_1_ = *D*_2_ = *D*^∗^ is a CSS or an evolutionary saddle point. In order to distinguish between these two possibilities, we inspect

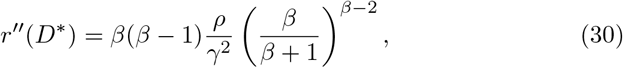

to find that *D*^∗^ is a CSS in case *β <* 1 (figure 5a) and an evolutionary saddle point if *β >* 1 (figure 5b).

**Figure 5.**
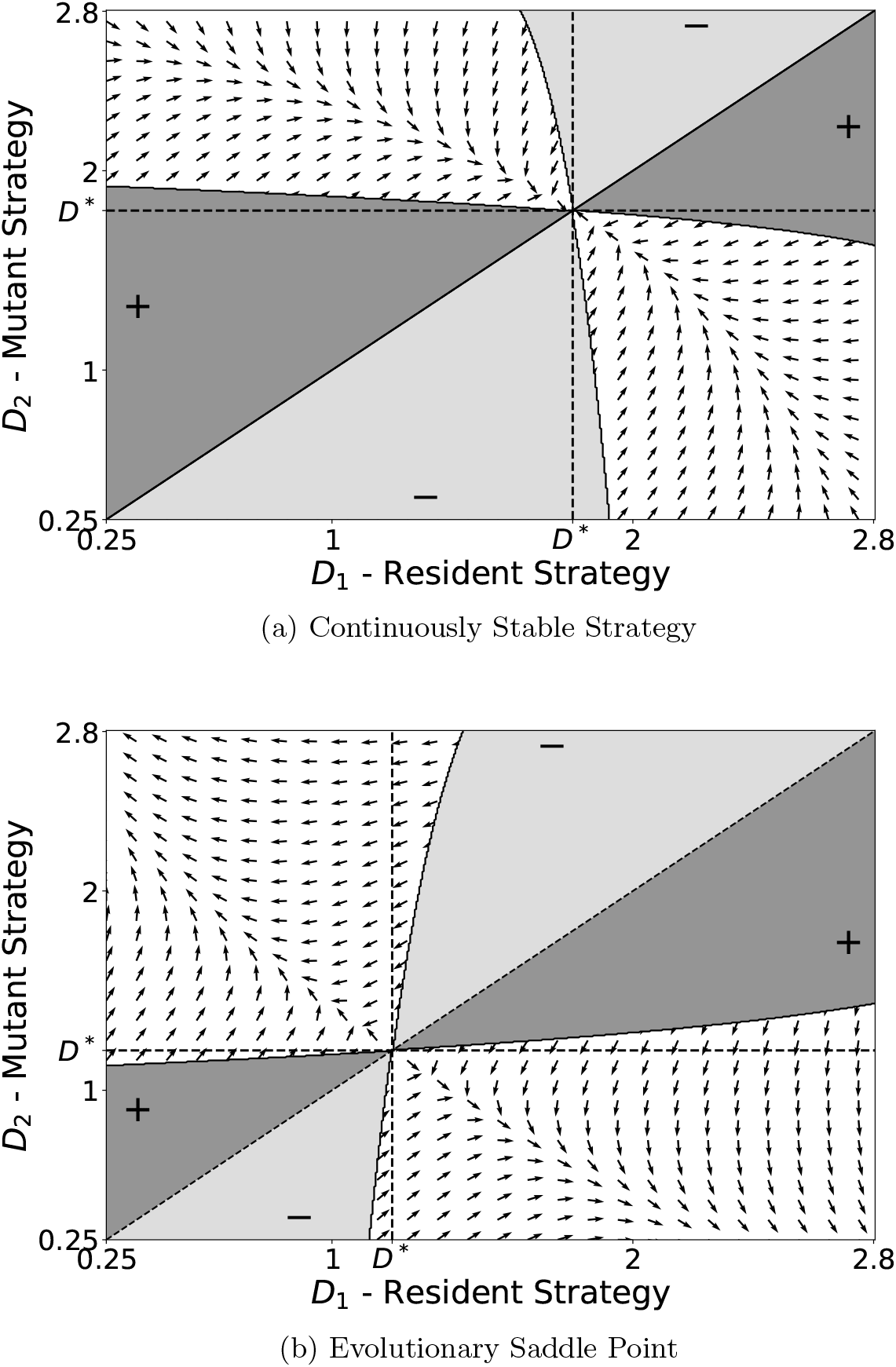
Pairwise invasibility plots for the imperfect clonal reproduction case, using *r*(*D*_*i*_) as defined in (27). *D*^∗^ is a CSS for *β* = 2*/*3 (5a) and an evolutionary saddle point for *β* = 3*/*2 (5b). Dimorphic pairs are possible in white regions, while arrows indicate the direction of speed increase. Gray regions are as in figure 4. Parameters used are *ρ* = *γ* = 3.

In case *β* = 1, the trade-off becomes a straight decreasing line, and the singular strategy *D*^∗^ = *γ/*2 is a monomorphic attractor. However, all dimorphic pairs with strategies *D*_1_ and *D*_2_ equidistant from *D*^∗^, that is, 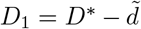 and 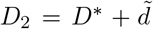, display precisely the same speed as the optimal single morph, i.e., 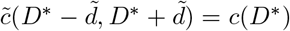. Hence, the test is inconclusive, and either two strategies equidistant from *D*^∗^ or a single strategy, *D*^∗^, itself, could be selected for at the leading edge.

Beyond the light and dark gray regions, in figures 5a and 5b, we have white regions, where dimorphic pairs are possible. These regions are filled with arrows pointing in the direction of dimorphic speed increase. In figure 5a, the arrows point to *D*^∗^, implying dimorphic populations would converge to a monomorphic population at the leading edge. The arrangement of the light and dark gray regions point to *D*^∗^ being attained monomorphically as well. Note also that the vertical (resp. horizontal) line through *D*^∗^ lies entirely inside the light (resp. dark) gray region, meaning that once the monomorphism is established, it is stable. In 5b, the arrows inside the white region point away from *D*^∗^, implying that it is repelling dimorphic populations, while the arrangement of the light and dark gray regions implies that it is approached monomorphically. Both the vertical and horizontal lines lie inside the white region, and therefore *D*^∗^, although approached monomorphically, is unstable and allow for dimorphisms to establish at the leading edge. The horizontal (resp. vertical) line also represents that mutants (resp. resident) with phenotype *D*_2_ = *D*^∗^ (resp. *D*_1_ = *D*^∗^) form a valid dimorphic pair with whichever resident (resp. mutant) phenotype *D*_1_ (resp. *D*_2_), or, in mathematical terms, 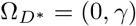 and 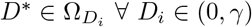.

### 3.2 Evolutionary Repellers

Now, we consider a trade-off relation in which every phenotype has a basal reproduction rate ρ, but the least dispersive phenotypes have a significant increase in their growth rates. Mathematically, we write

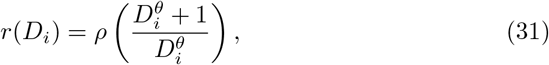

with *D*_*i*_ ∈ (0, *∞*) and *θ*, ρ > 0 as shape and scale parameters, respectively. This trade-off is concave-up for any parameter choice, and *θ* sets how strongly growth rates and diffusion coefficients are anti-correlated.

The singular strategy, *D*^∗^, is given by

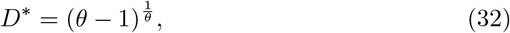

which is only positive if *θ >* 1. In the case *θ <* 1, the monomorphic speed *c*(*D*_*i*_) is monotonically increasing, and our analysis implies larger *D*_*i*_ is selected for at leading edge.

The second derivative of the monomorphic speed at *D*_2_ = *D*^∗^ is

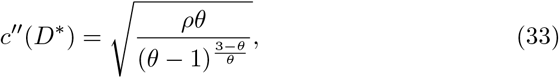

and therefore, *D*^∗^ is an evolutionary repeller both in the perfect and imperfect clonal reproduction cases for *θ >* 1. In figure 6a, we have the pairwise invasibility plot for perfect clonal reproduction. Note that resident strategies *D*_1_ *< D*^∗^ are invaded by mutant strategies *D*_2_ *< D*_1_, represented by the light (resp. dark) gray coloring and the minus (resp. plus) sign above (resp. below) the line *D*_1_ = *D*_2_. On the other hand, resident strategies *D*_1_ *> D*^∗^ are invaded by strategies *D*_2_ *> D*_1_, represented graphically in similar fashion. In the case of imperfect clonal reproduction, the arrows inside the white region in figure 6b are pointing away from *D*^∗^, showing that both initially monomorphic and dimorphic populations are expected to evolve phenotypes away from *D*^∗^ at the leading edge.

**Figure 6.**
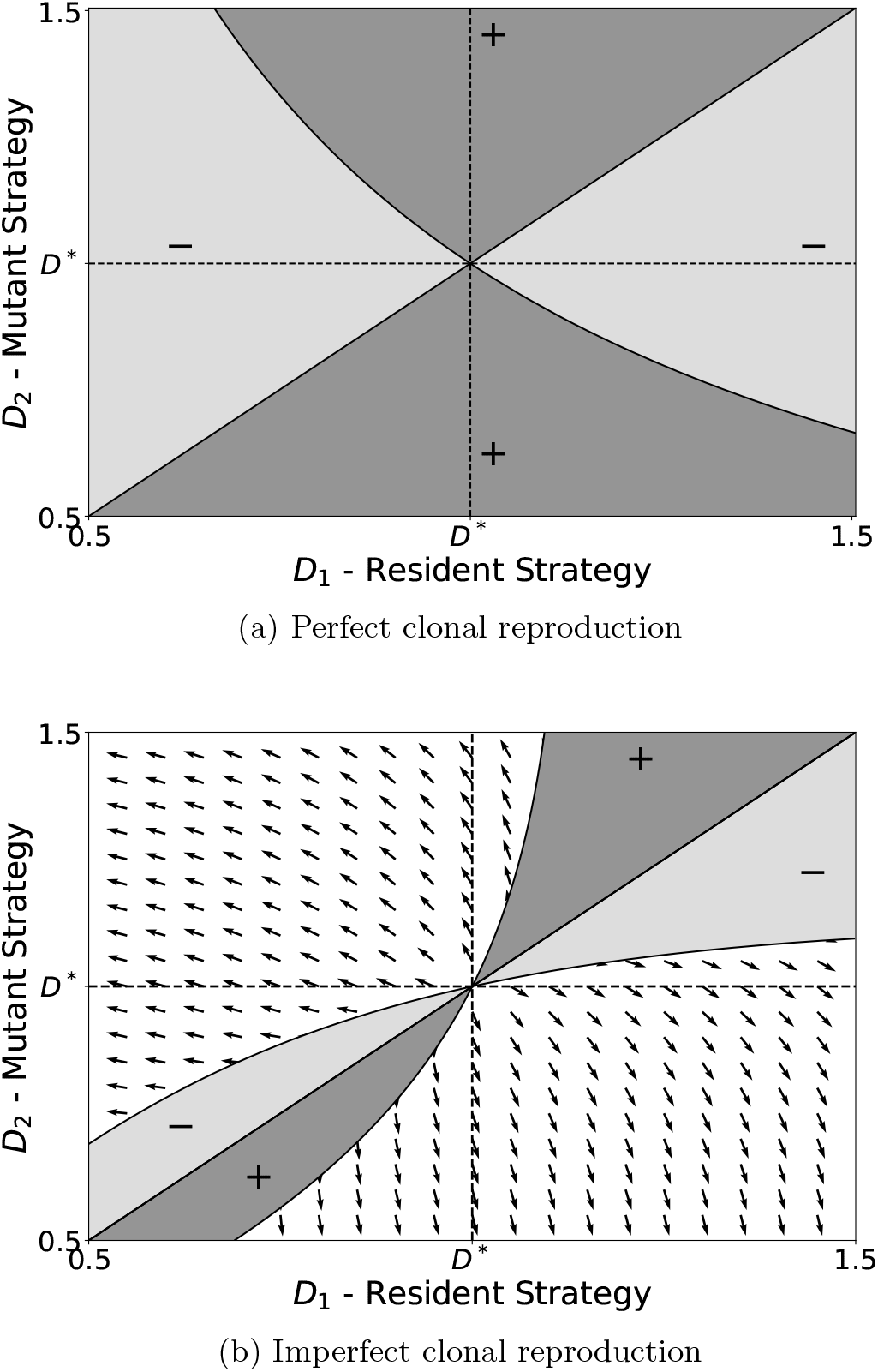
Pairwise invasibility plots for both perfect and imperfect clonal reproduction cases, using *r*(*D*_*i*_) as defined in (31). White, light and dark gray regions have the same meaning as in figure 5a. Note that *D*^∗^ is a repeller in both cases, as mutant strategies away from it invade resident strategies closer to it. Parameters are *ρ* = 3 and *θ* = 2.

## 4 Discussion

We have inspected how evolution shapes the leading edge of biological invasions in the context of adaptive dynamics. We found conditions for a mutant population to establish at the leading edge considering different phenotypic inheritance mechanisms. Such conditions depend on mutant’s asymptotic spreading speed being higher than resident’s asymptotic spreading speed. When offspring are identical to their parents, the fastest monomorphic population is selected fo at the leading edge. On the other hand, if inheritance mechanisms allow mutant offspring to inherit resident phenotypes and vice versa, then dimorphisms can be maintained and favored at the leading edge.

The notion of spreading speed being maximized at the leading edge is not exclusive to our work. Phillips and Perkins [2019], following a similar modeling framework for natural selection present in Otto and Day [2007] (ch. 3), showed that if fitness has both a spatial and a temporal component, then the species might develop phenotypes that do not maximize fitness in the classical sense (i.e. temporal component). Instead, phenotypes with an optimal combination of both spatial and temporal components evolve. The spatial component can be associated with dispersal phenotypes, while the temporal component can be associated with demographic or reproduction phenotypes. In the scope of F-KPP type reaction-diffusion equations, this translates into the species maximizing the product of intrinsic growth rates and diffusion coefficients in the form of speed, i.e., 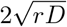, as described here and in Deforet et al. [2019].

However, as we illustrated in this work, under trade-offs between reproduction and dispersal, maximizing growth and dispersal can lead to different evolutionary outcomes (3). In the case of perfect clonal reproduction, either an optimal strategy can be achieved through subsequent mutations (figure 4), or individuals of the species can invest fully in reproduction or dispersal (figure 6a). In both cases, the population remains monomorphic at the leading edge, and an increase in spreading speed, due to such evolutionary process, leads to an accelerating invasion, as it was observed, for example, in the cane toad invasion of Australia [Phillips et al., 2006].

Trade-offs also play a significant role in the viability of dimorphisms. Our results in these cases are an extension of the main results in Keenan and Cornell [2021] and Poloni and Lutscher [2023]. In the case of concave-down trade-offs, their results show that polymorphic populations ^2^ cannot spread faster than their single fastest morph, and that this morph is the most abundant at the leading edge. Here we recover the result, instead, by showing that concavedown trade-offs lead to continuously stable strategies, which are approached monomorphically and dimorphically, through the course of the biological invasions (figure 5b). In the case of concave-up trade-offs, previous results showed that polymorphic populations can display anomalous spreading speeds, and that said speed and leading edge trait distributions depend heavily on the extremes of the trait spectrum. This, in turn, is recovered here by showing that evolution can take two distinct routes. Dimorphisms are either selected for in the case of an evolutionary saddle point (figure 5a), or simply remain in the population if present on the early stages of invasion in the case of an evolutionary repeller (figure 6b).

Noticeably, in the presence of a continuously stable strategy, evolution pushes initially dimorphic populations to become monomorphic at the leading edge. This in turn implies that, at least transiently, mechanisms that allow for the maintenance of two or more morphs in a population might be lost in the course of an invasion. On the other hand, in the presence of an evolutionary saddle point, species could be pushed to develop such mechanisms, in order to gain colonization advantages over an optimal single morph. This might be particularly relevant for different winged insect taxa, since many species, such as aphids, wasps and crickets can display wing dimorphisms relying on epigenetic factors Roff [1986], Mole and Zera [1993], Cônsoli and Vinson [2002], Braendle et al. [2006], and range expansions can serve as a driver for such evolutionary phenomena to take place in novel species.

The criteria for invasion at the leading edge found here was solely based on the linearized analysis. A more precise investigation of general conditions for which a small density of mutant population can take over the leading edge is a novel mathematical challenge. Formally, it leads to an eigenvalue problem that may be addressed following and extending existing results in Berestycki and Rossi [2015], Girardin and Lam [2019], Liu et al. [2021]. Our numerical investigations (supplemental material) indicate that the criteria used here are valid at least when both resident and mutant populations are small, i.e., at the leading edge of the wavefront. However, mutant populations initially closer to the front might be washed away by resident population range expansion, and therefore never establish, despite displaying the highest asymptotic spreading speed.

The framework presented here can be useful in the analysis of different range expansion phenomena that present a linearly determinate speed, i.e., when small population densities are able to grow at newly colonized regions. For example, populations spreading in heterogeneous landscapes can present linearly determinate speeds [Yurk and Cobbold, 2018, Cobbold et al., 2022, Hamel et al., 2022], and the framework could be used to study, for example, habitat preference evolution throughout range expansion. Predator invasions upon established resident prey also display linearly determinate speeds Petrovskii and Malchow [2000], Malchow [1997], where evolutionary phenomena may take place, driving predator phenotypes at the leading edge in different directions than expected at the core of invasion. Furthermore, although the analysis is entirely presented here in terms of reaction-diffusion equations, the equivalent framework in terms of integro-difference equations can also be used to explore, for example, the evolution of different movement strategies other than Brownian movement [Lutscher, 2019].

## 5 Acknowledgments

We would like to thank King-Yeung (Adrian) Lam for discussions regarding the eigenvalue problem defining the invasion exponent, and Vincent Calvez for pointing out important references and discussion.

## Footnotes

1 given it is initially concentrated in some region of space, i.e., a compactly supported initial condition

2 consisting of more than two morphs

