## Supplemental Material for "Evolutionary dynamics at the leading edge of biological invasions"

#### 1 Mathematical Analysis of the Imperfect Clonal Reproduction case

We start by re-stating the model

$$\begin{cases} \partial_t u_1 = D_1 \partial_x^2 u_1 + r_1 u_1 (1 - u_1 - a_{uv}(v_1 + v_2)), \\ \partial_t v_1 = D_1 \partial_x^2 v_1 + (r_1(1-\alpha_{21}) v_1 + r_2 \alpha_{12} v_2)(1 - v_1 - v_2 - a_{vu} u) \\ \partial_t v_2 = D_2 \partial_x^2 v_2 + (r_2(1-\alpha_{12}) v_2 + r_1 \alpha_{21} v_1)(1 - v_2 - v_1 - a_{vu} u), \end{cases} \tag{1}$$

and linearizing at the leading edge, where both mutants and residents are rare, we get

$$\begin{cases} \partial_t u_1 = D_1 \partial_x^2 u_1 + r_1 u_1, \\ \partial_t v_1 = D_1 \partial_x^2 v_1 + r_1 v_1 + r_2 \alpha_{12} v_2 - r_1 \alpha_{21} v_1, \\ \partial_t v_2 = D_2 \partial_x^2 v_2 + r_2 v_2 + r_1 \alpha_{21} v_1 - r_2 \alpha_{12} v_2. \end{cases} \tag{2}$$

For mathematical simplicity, we assume $\mu = r_1\alpha_{21} = r_2\alpha_{12}$. The symmetry in $r_j\alpha_{ij}$ has little impact on the results shown here, as long as $r_2\alpha_{12}$ and $r_1\alpha_{21}$ are of the same order [Elliott and Cornell, 2012].

Assuming a traveling front of the form $u_1 = \phi_1 \mathrm{e}^{-\sigma(x-ct)}$, $v_i = \psi_i \mathrm{e}^{-\sigma(x-ct)}$, and plugging it into the linearized equation we get

$$\Lambda(\sigma,\mu) \begin{bmatrix} \phi_1 \\ \psi_1 \\ \psi_2 \end{bmatrix} = \begin{bmatrix} w_1(\sigma) & 0 & 0 \\ 0 & w_1(\sigma) - \mu/\sigma & \mu/\sigma \\ 0 & \mu/\sigma & w_2(\sigma) - \mu/\sigma \end{bmatrix} \begin{bmatrix} \phi_1 \\ \psi_1 \\ \psi_2 \end{bmatrix}. \tag{3}$$

which is an eigenvalue problem for $\Lambda(\sigma,\mu)$, with $w_i(\sigma) = r_i/\sigma + D_i\sigma$. Note that $c_i = c(D_i) = \inf_{\sigma>0} w_i(\sigma)$ for $c$ as defined in the main text. Also note that $w_1(\sigma)$ is an eigenvalue of the matrix, while the other two eigenvalues are given by solving

$$w(\sigma,\mu)\begin{bmatrix}\psi_1\\ \psi_2\end{bmatrix} = \begin{bmatrix} w_1(\sigma)-\mu/\sigma & \mu/\sigma \\ \mu/\sigma & w_2(\sigma)-\mu/\sigma\end{bmatrix}\begin{bmatrix}\psi_1\\ \psi_2\end{bmatrix}, \tag{4}$$

$$w(\sigma,\mu)\mathbf{\Psi}(\sigma,\mu) = \mathbf{A}(\sigma,\mu)\mathbf{\Psi}(\sigma,\mu), \tag{5}$$

where $\mathbf{\Psi}(\sigma,\mu) = (\psi_1,\psi_2)$, with $\psi_i \equiv \psi_i(\sigma,\mu)$, and $\mathbf{A}(\sigma,\mu)$ the 2 by 2 matrix.

The eigenvalues, $w(\sigma,\mu)$, are

$$w(\sigma,\mu) = \frac{1}{2}\left\{w_1(\sigma)+w_2(\sigma)-2\mu/\sigma \pm \sqrt{(w_1(\sigma)-w_2(\sigma))^2+4\mu^2/\sigma^2}\right\}, \tag{6}$$

from where we readily see that in the limit $0<\mu\ll \min_i(r_i)$, we have that the dominant eigenvalue for each value of $\sigma$, $w(\sigma,\mu\approx 0)$, is

$$w(\sigma,\mu\approx 0) = \max_i\{w_i(\sigma)\} - \frac{\mu}{\sigma} + O(\mu^2), \tag{7}$$

and for the sake of brevity in notation, we will refer to $w(\sigma,\mu\approx 0)$ just as $w(\sigma)$ from this point.

Assuming the linear determinacy of the asymptotic spreading speed, $\Lambda \equiv \Lambda(\mu) = \inf_{\sigma>0}\Lambda(\sigma,\mu)$ we have that it must be given by

$$\Lambda(\mu) = \max(c_1, \inf_{\sigma>0} w(\sigma,\mu)). \tag{8}$$

Note that the species spreading speed is given by the largest asymptotic spreading speed available between resident and mutant populations. Since $w(\sigma,\mu)$ depends on both $w_i(\sigma)$, it is helpful to plot the curves $w_1(\sigma)$, $w_2(\sigma)$ and $w(\sigma,\mu)$ in the limit $0<\mu\ll\min_i(r_i)$, as performed in figure 1, to realize that such curves can only be plotted in two qualitatively distinct ways.

The plots in figures 1a show us that one of these qualitatively distinct scenarios have $\inf w(\sigma)\approx c_i = 2\sqrt{r_iD_i}$, with the infimum of $w(\sigma)$ taking place at $\sigma\approx\sigma_i^* = \sqrt{r_i/D_i}$, i.e., the mutants will travel with the speed of its fastest single morph. The second case, displayed in plot 1b, is found when the infimum takes place at the intersection between $w_1(\sigma)$ and $w_2(\sigma)$, namely, at $\sigma\approx\sigma_d^* = \sqrt{(r_1-r_2)/(D_2-D_1)}$, for which we have $\inf_{\sigma>0} w(\sigma)\approx\tilde{c} = \sqrt{(r_1+r_2)(D_1+D_2)} > \max_i(c_i)$, i.e., the anomalous spreading speed.

Note, however, that the asymptotic spreading speed, and by consequence, the leading edge morph distributions, are not necessary continuous at $\mu=0$ when an anomalous spreading speed is possible for $\mu>0$. Take, for example, the case where morph 2 is the fastest, then, for $\mu=0$, the speed would be $\Lambda(0)=c_2$, and the leading edge morph distribution would be the corresponding eigenvector to $w(\sigma_2^*,0)=c_2$ in (4), given by $\mathbf{\Psi}(\sigma_2^*,0)=(\psi_1,\psi_2)=(0,1)^T$.

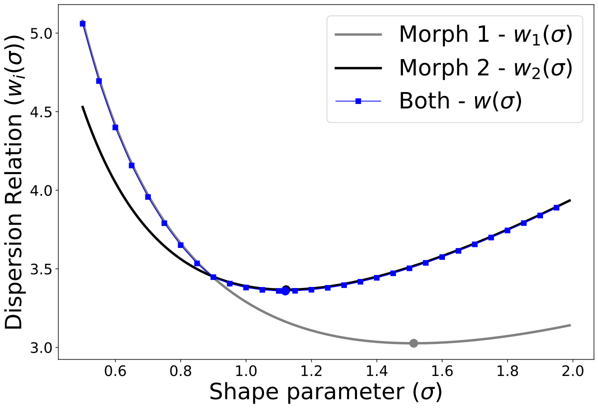


(a) Infimum of $w(\sigma)$ on Morph 2 line

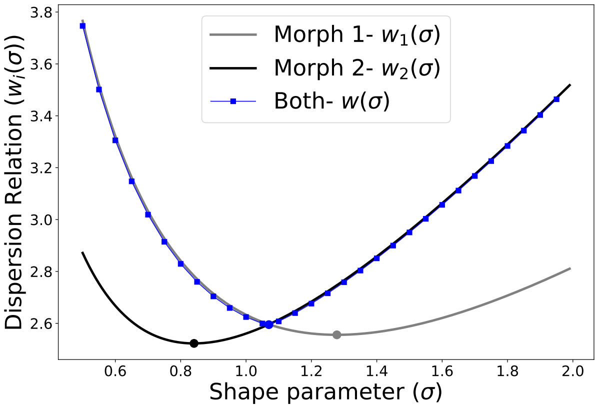


(b) Infimum of $w(\sigma)$ lies at the intersection

Figure 1: Dispersion relations for isolated morphs in black and gray, and with imperfect clonal reproduction in blue line with squares. The black and gray dots are the asymptotic spreading speeds, the minimum of dispersion relations w.r.t. $\sigma$. Note how on 1a the blue dot superposes the black dot, both lie on the blue line with squares, and the speed of both morphs together is the same as morph 2. While in 1b the blue dot lies at the intersection between $w_1(\sigma)$ and $w_2(\sigma)$, and the asymptotic spreading speed of both morphs together is higher than in isolation (the anomalous spreading speed)

Now, increase $\mu$ by some small amount, and the asymptotic spreading speed jumps from $c_2$ to $\Lambda(\mu \approx 0) \approx \tilde{c}$, with $\tilde{c} - c_2$ non-infinitesimal in general, the corresponding eigenvector to $\tilde{c}$ may also change abruptly from $(0, 1)$ to a different vector.

In the case anomalous spreading speeds are not present, the asymptotic spreading speed is continuous close to $\mu = 0$. Since for $\mu = 0$ we have $\Lambda(0) = \max_i c_i$, while for $\mu$ small, we have $w(\sigma) \approx \max_i(w_i(\sigma)) + \mu/\sigma^2$ with infimum at $\sigma = \sigma_i^* + \mu/c_i$, implying $\Lambda(\mu \approx 0) = c_i - \mu h(D_i, r_i)$, with $h$ a positive and bounded function for $D_i, r_i > 0$. Hence, $\Lambda(\mu \approx 0) - \Lambda(0) = -\mu h(r_i, D_i)$, with $i$ representing the fastest morph. This implies that, when the fastest mutant morphs are those displaying the same phenotypes as the resident population and no anomalous spreading speeds are possible, residents are not invaded by mutants, since displaying both novel and resident phenotypes slows down their spreading speed, i.e., $\Lambda(\mu \approx 0) < \Lambda(0)$ [1].

In the main text we claim that mutants displaying both novel and resident phenotypes will become monomorphic, despite $\mu > 0$, if no anomalous spreading speeds are formed and given $c_2 > c_1$, i.e., the novel set of phenotypes provides a faster asymptotic spreading speed. As aforementioned, if $c_2 > c_1$ and $\mu = 0$, the dominant eigenvector of (4) is $\mathbf{\Psi}_1(\mu = 0) = (0, 1)^T$, while the other eigenvector is $\mathbf{\Psi}_2(\mu = 0) = (1, 0)^T$ (we omit the notation, but both are taken at $\sigma = \sigma_2^*$). We exploit this to write

$$\mathbf{\Psi}_1(\mu) \approx \mathbf{\Psi}_1(0) + \mu \partial_\mu \mathbf{\Psi}_1(0), \quad \mu \approx 0. \tag{9}$$

Calculating $\partial_\mu \mathbf{\Psi}_1(0)$ requires some care, but following Caswell [2001], one finds

$$\begin{aligned}
\partial_\mu \mathbf{\Psi}_1(0) &= \frac{1}{c_2 - w_1(\sigma_2^*)} (\partial_\mu \mathbf{A} \mathbf{\Psi}_1(0) \cdot \mathbf{\Psi}_2(0)) \mathbf{\Psi}_2(0), && (10)\\
&= \frac{1}{\sigma_2^*(c_2 - w_1(\sigma_2^*))} \mathbf{\Psi}_2(0), && (11)\\
&= -\frac{1}{r_2\left(\frac{r_1}{r_2} + \frac{D_1}{D_2} - 2\right)} \mathbf{\Psi}_2(0), && (12)
\end{aligned}$$

Therefore,

$$\mathbf{\Psi}_1(\mu) \approx \mathbf{\Psi}_1(0) - \frac{\mu}{r_2\left(\frac{r_1}{r_2} + \frac{D_1}{D_2} - 2\right)} \mathbf{\Psi}_2(0), \tag{13}$$

from where we conclude that the shift from the monomorphic population $\mathbf{\Psi}_1(0)$ to $\mathbf{\Psi}_1(\mu)$ should be of order $\mu/r_2 \ll 1$, as long as $r_1/r_2 + D_1/D_2 < 2$, hence, the population is mostly monomorphic with phenotypes $(r_2, D_2)$. However, if $r_1/r_2 + D_1/D_2 > 2$, $\mathbf{\Psi}_1(\mu)$ is no longer positive, and hence it can't correspond to a dominant eigenvalue, since matrix $\mathbf{A}$ is positive and Perron-Frobenius theorem applies. Therefore $c_2$ is no longer the asymptotic spreading speed of the population.

[1] The result can be stated more generally, see Morris et al. [2019], Poloni and Lutscher [2023]

We can use the same argument for monomorphic fronts of speed $c_1$, to find that the general condition for $c_i$ to be the assymptotic spreading speed of the mutant population is

$$\frac{r_j}{r_i} + \frac{D_j}{D_i} - 2 < 0,\ i, j = 1, 2, i \neq j. \tag{14}$$

When conditions (14) are not met, neither $c_1$ or $c_2$ can be the asymptotic spreading speeds. To show that such cases lead to anomalous asymptotic spreading speeds, we follow the analysis from Elliott and Cornell [2012]. Let $\tilde{\sigma}^2 = {\sigma_d^*}^2 + \epsilon$, and for $\mu$ small, we should have that $\tilde{\sigma}$ is a critical point of $w(\sigma)$ with $\epsilon$ also small, i.e.,

$$\partial_s w(\sigma = \tilde{\sigma}) = 0, \tag{15}$$

such that $w$ reaches its infimum at $\tilde{\sigma}$. This leads to the following equation

$$((D_\delta \tilde{\sigma}^2 + r_\delta)^2 + 4\mu^2)(D_\sigma \tilde{\sigma}^2 - r_\sigma - 2\mu)^2 = (D_\delta^2 \tilde{\sigma}^4 - r_\delta^2 - 4\mu^2)^2, \tag{16}$$

where $D_\delta = D_1 - D_2$ and $D_\sigma = D_1 + D_2$ and similarly for $r_\delta$ and $r_\sigma$.

Substituting the expression for $\tilde{\sigma}^2$ in (16), and noticing that $D_\delta {\sigma_d^*}^2 = -r_\delta$, $D_\delta^2 {\sigma_d^*}^4 = r_\delta^2$, $r_\sigma D_\delta + r_\delta D_\sigma = 2(r_1 D_1 + r_2 D_2)$, and ignoring terms of $\mu^p \epsilon^q$ with order $p + q \geq 3$, we get

$$\frac{4}{D_\delta^2}\left(\frac{\mu}{\epsilon}\right)^2 = \frac{r_\delta^2 D_\delta^2}{(r_2 D_2 - r_1 D_1)^2} - 1. \tag{17}$$

Since the left side of (17) is positive, we have that

$$\frac{r_\delta^2 D_\delta^2}{(r_2 D_2 - r_1 D_1)^2} \geq 1 \tag{18}$$

which is an alternative way to display the compliment of conditions (14) in $(D_1, D_2, r_1, r_2) \in \mathbb{R}_+^4$.

Now, for the leading edge morph distribution, we use the fact that at $\sigma = \sigma_d^*$, $\tilde{c} = \sqrt{(r_1 + r_2)(D_1 + D_2)} = c_1(\sigma_d^*) = c_2(\sigma_d^*)$,therefore, at this particular point

$$\boldsymbol{A}(\sigma_d^*, \mu) = \begin{bmatrix} \tilde{c} - \mu/\sigma_d^* & \mu/\sigma_d^* \\ \mu/\sigma_d^* & \tilde{c} - \mu/\sigma_d^*, \end{bmatrix} \tag{19}$$

which has dominant eigenvalue $w(\sigma_d^*, \mu) = \tilde{c}$ and corresponding eigenvector $\boldsymbol{\Psi}_1(\sigma_d^*, \mu) = (1/\sqrt{2}, 1/\sqrt{2})$. The other eigenpair is $w(\sigma_d^*, \mu) = \tilde{c} - 2\mu/\sigma_d^*$ with $\boldsymbol{\Psi}_2(\sigma_d^*, \mu) = (-1/\sqrt{2}, 1/\sqrt{2})$ . Note, however, that $w(\sigma_d^*, \mu) \neq \inf_{\sigma>0} w(\sigma, \mu)$, but as we saw before, if $\mu$ is small, then the infimum takes place at $\tilde{\sigma} = \sigma_d^* + \epsilon$, with $\epsilon$ small. A similar calculation as performed for equation (9) can be carried out here, revealing

$$\boldsymbol{\Psi}_1(\sigma_d^* + \epsilon, \mu) \approx \boldsymbol{\Psi}_1(\sigma_d^*, \mu) + \frac{\epsilon \sigma_d^*}{2\mu} \frac{(r_\delta D_\delta - r_\delta D_\delta)}{r_\delta} \boldsymbol{\Psi}_2(\sigma_d^*, \mu), \quad \epsilon, \mu \approx 0. \tag{20}$$

Therefore

$$\mathbf{\Psi}_1(\sigma_d^* + \epsilon, \mu) \approx \mathbf{\Psi}_1(\sigma_d^*, \mu), \tag{21}$$

and the population is strongly dimorphic, as pointed out in the main text. Because the first order approximation term yields zero, the logical step would be to calculate the second order term. However, the second order term is hard to gain any insight and would likely depend on $\epsilon^2/\mu^2$, leading us back to relation (17).

### 2 Review of Adaptive Dynamics

We provide this quick review on adaptive dynamics for completeness, but also to illustrate how the sources for dimorphism found in the main paper are an emergent property of range expansion processes, and possibly other spatial contexts, but unlikely to appear in classical adaptive dynamics. We still refer to Diekmann [2003] for a complete guide on adaptive dynamics.

In adaptive dynamics, we assume that a resident population, $u_1$, with phenotype $z_1$ grows and establishes to $u_1 = u^*(z_1)$ at ecological timescales. On doing so, the resident population sets the environment in the equilibrium $E_1 = E(z_1)$. After a longer time, a rare/small mutant population, $u_2$, with phenotype $z_2$, appears, and needs to grow in such environment. An ordinary differential equation description of how mutant population changes in time, $t$, is given by

$$\frac{\mathrm{d}u_2}{\mathrm{d}t} = G(u_2, u_1, z_2, E). \tag{22}$$

Where $G$ is a growth function, with $G(u_2 = 0, \cdot) = 0$ (if there is no mutant population, there are no changes in mutant population size). Since we assume that mutant population is rare, i.e., $u_2 \approx 0$, resident population is established at $u_1 = u^*(z_1)$, and that the environment is established at $E_1 = E(z_1)$, we can write

$$G \approx G(0, u^*(z_1), z_2, E_1(z_1)) + u_2 \partial_{u_2} G(0, u^*(z_1), z_2, E_1(z_1)) = u_2 s(z_1, z_2), \tag{23}$$

where we defined $s(z_1, z_2) = \partial_{u_2} G(0, u^*(z_1), z_2, E_1(z_1))$.

Now, plugging (23) back to equation (22) we find

$$\frac{\mathrm{d}u_2}{\mathrm{d}t} = u_2 s(z_1, z_2), \tag{24}$$

and find that $s = s(z_1, z_2)$ is the growth rate of mutants with phenotype $z_2$ in an environment set by residents of phenotype $z_1$, i.e., $s$ is the invasion exponent. It follows that $s > 0$ leads to successful invasion of mutants (and failed invasions otherwise). These mutants will then grow and establish, at population levels $u_2 = u^*(z_2)$, while setting the environment to $E_2 = E(z_2)$. From here, the analysis of a novel mutant, $u_3$, can be pursued in a similar manner.

Note that $z_2$ (mutant's phenotype) is expected to be close to $z_1$ (resident phenotype), so that

$$s(z_1, z_2) \approx s(z_1, z_1) + (z_2 - z_1)\partial_{z_2} s|_{z_2=z_1} + \frac{(z_2 - z_1)^2}{2}\partial^2_{z_2} s|_{z_2=z_1}. \tag{25}$$

By definition we have $s(z_1, z_1) = 0$, i.e., a mutant with the same phenotype of the resident population can neither grow or decay, since it would face the environment set by residents of phenotype $z_1$ itself, which is at equilibrium.

The main goal is finding critical values of phenotypes/strategies which lead to, for example, a halt in evolution, and if such points can be achieved via subsequent mutant invasions. For that, we deploy the selection gradient, $\partial_{z_2} s|_{z_2=z_1}$, which indicates the direction of increase in $s$. When such test fails, i.e., $\partial_{z_2} s = 0$ at $z_1 = z_2 = z^*$, we have a singular strategy, $z^*$. We then inspect the second order derivatives of $s$ at $z_1 = z_2 = z^*$ to study the properties of such singular strategy. The most relevant points are summarized in table 1.

| Derivative signs | Qualitative Behavior of singular strategy |
|---|---|
| $s_{22} < 0$ | Evolutionary Stable Strategy (ESS) |
| $s_{11} > s_{22}$ | Convergence Stable |
| $s_{22} < 0,\ s_{11} > s_{22}$ | Continuously Stable Strategy (CSS) |
| $s_{22} < 0,\ s_{22} > -s_{11}$ | Dimorphisms near the singular strategy |
| $s_{22} > 0,\ s_{11} > s_{22}$ | Branching Point |
| $s_{22} > 0,\ s_{11} < s_{22}$ | Evolutionary Repeller |

Table 1: Summary of main qualitative behavior of singluar strategies. We define $s_{ij} = \partial^2_{z_i,z_j} s|_{z_2=z_1=z^*}$.

We can write the analogous model for a mutant population with two distinct strategies in classical adaptive dynamics framework. Let the mutant populations $v_1$ and $v_2$, display strategies $z_{v1}$ and $z_{v_2}$, respectively. In an environment set by a resident population with strategy $z_{u1}$, the equations for mutant growth are

$$\begin{cases} \frac{\mathrm{d}v_1}{\mathrm{d}t} &= v_1 s(z_{u1}, z_{v1}) + \mu(v_2 - v_1) \\ \frac{\mathrm{d}v_2}{\mathrm{d}t} &= v_2 s(z_{u1}, z_{v2}) + \mu(v_1 - v_2) \end{cases} \tag{26}$$

The invasion coefficient of the mutant population is given by $\tilde{s}(z_{u1}, z_{v1}, z_{v2})$, defined as the dominant eigenvalue of matrix

$$\mathbf{A} = \begin{bmatrix} s(z_{u1}, z_{v1}) - \mu & \mu \\ \mu & s(z_{u1}, z_{v2}) - \mu \end{bmatrix}. \tag{27}$$

To have a saddle point in strategy $z^*_u$, we must have that $s(z^*_u, z_v) < 0$ for all $z_v \neq z^*_u$, while $\tilde{s}(z^*_u, z_{v1}, z_{v2}) > 0$ for some $z_{v1}$ and $z_{v2}$ close to $z^*_u$. Clearly, in this formulation, a saddle point is non feasible since $\mathrm{tr}\mathbf{A} < 0$ and $\det \mathbf{A} > 0$, leading to all eigenvalues of $\mathbf{A}$ being negative, and hence $\tilde{s}(z^*_u, z_{v1}, z_{v2}) < 0$.

Introducing new growth rates $G_i(z_{u1}, z_{v1}, z_{v2})$, $i = 1, 2$, such that mutant morph $i$ follows the equation

$$\frac{\mathrm{d}v_i}{\mathrm{d}t} = v_i G_i(z_{u1}, z_{v1}, z_{v2}) + \mu(v_j - v_i), \tag{28}$$

we explicitly assume that both morphs display a different growth rate together from that they display if alone. We then have matrix $A$ become

$$\mathbf{A} = \begin{bmatrix} G_1(z_{u1}, z_{v1}, z_{v2}) - \mu & \mu \\ \mu & G_2(z_{u1}, z_{v1}, z_{v2}) - \mu \end{bmatrix}. \tag{29}$$

For a singular strategy $z_u^*$ to be a saddle point, we must have that $G_i(z_u^*, z_{vi}, z_{vi}) = s(z_u^*, z_{vi})$ has a maximum at $z_{vi} = z_u^*$ and the dominant eigenvalue of $\mathbf{A}$ is positive in some neighborhood of said maximum.

### 3 Eigenvalue problem and precise invasion exponent definition

Let the monomorphic resident population density be at its traveling wave profile, i.e., $u_1 = u_1^*(z = x - c_1 t)$. Upon introducing a small density of a monomorphic population with phenotype $D_2$ at the leading edge of the resident population, we have, on the moving frame $z = x - c_1 t$, the eigenvalue problem

$$\lambda u_2 = D_2 u_2'' + c_1 u_2' + r_1(1 - u_1^*(z))u_2, \tag{30}$$

where $\cdot' = \partial_z \cdot$. Changing the variable $u_2$ for $\psi = u_2 \mathrm{e}^{-c_1 z/2D_2}$, we set the eigenvalue problem to

$$\lambda\psi = D_2\psi'' + r_2(1 - (c_1/c_2)^2 - u_1^*(z))\psi. \tag{31}$$

For $z \in [a, b] \subset \mathbb{R}$ and $\psi$ with Dirichlet boundary conditions, Krein-Rutman theorem grants that a dominant eigenvalue, $\lambda_1$, exists and has an unique correspondent eigenfunction of one sign. However, in this problem we are concerned on $z \in \mathbb{R}$, i.e., an unbounded domain, where a similar notion of dominant eigenvalue is not straightforward to obtain. Berestycki and Rossi [2015] provide the following notion of dominant eigenvalue for problems in unbounded domains:

$$\lambda_1 = \sup\{\lambda : \exists\psi \in C^2(\Omega), \psi > 0 \text{ and } (L - \lambda)\psi \leq 0 \text{ a.e. in } \Omega\}, \tag{32}$$

where $L\psi$ yields the right-hand side of equation (31) and $\Omega$ is the domain (here $\Omega = \mathbb{R}$). This notion grants some of the desired properties for the dominant eigenvalue of (31), given the settings for our problem ($u_1^*(z)$ is continuous, monotone and bounded in $[0, 1]$, $D_2 > 0$ and $c_i \in \mathbb{R}$). That is, there exists a dominant eigenvalue such that $\lambda_1 = \sup(E)$, $E$ the set of eigenvalues, and its corresponding eigenfunction is unique and of one sign (almost everywhere). Our invasion coefficient in the main text, $s_{D_1}(D_2)$, is precisely the dominant eigenvalue, $\lambda_1$.

Because $0 \leq u_1^*(z) \leq 1$, we have $L_1\psi \leq L\psi \leq L_0\psi$ ($L_0$ and $L_1$ the operator $L$ with $u_1^*$ replaced by 0 and 1, respectively), it follows that $\lambda_1$ is also bounded, i.e.,

$$\lambda_1 \in \left[ -r_2 \left( \frac{c_1}{c_2} \right)^2 ,\ r_2 \left( 1 - \left( \frac{c_1}{c_2} \right)^2 \right) \right]. \tag{33}$$

The lower bound can be interpreted as mutant introductions to regions where resident population is well established, $u_1^*(z \to -\infty) = 1$, and the upper bound corresponding to regions far-off the leading edge, $u_1^*(z \to -\infty) = 0$. The later is the argument used in Deforet et al. [2019], i.e., if we are considering introductions at the leading edge, the dominant eigenvalue must be closer to its upper bound, and hence $\text{sign}\{\lambda_1\} = \text{sign}\{c_2 - c_1\}$. It is clear that for $c_2 < c_1$ invasions are not possible.

Note, nonetheless, that the problem gets more intricate for a mutant with phenotypic plasticity. Let us say with phenotypes $D_1$ and $D_2$, and corresponding densities $v_1$ and $v_2$. The eigenvalue problem reads

$$\begin{aligned} \lambda v_1 &= D_1 v_1'' + c_1 v_1' + \rho_1 (1 - u_1^*(z)) u_1 + \mu(v_2 - v_1), \\ \lambda v_2 &= D_2 v_2'' + c_1 v_2' + \rho_2 (1 - u_1^*(z)) v_2 + \mu(v_1 - v_2). \end{aligned} \tag{34}$$

Developing a notion of dominant eigenvalue for such problems and estimating them is a possible venue of future research.

### 4 Hessian matrix of $\tilde{c}$

The Hessian matrix of $\tilde{c}$ is

$$\mathbf{H}_{\tilde{c}}(D_1, D_2) = \begin{bmatrix} \partial^2_{D_1} \tilde{c} & \partial^2_{D_1, D_2} \tilde{c} \\ \partial^2_{D_2, D_1} \tilde{c} & \partial^2_{D_2} \tilde{c} \end{bmatrix}. \tag{35}$$

We are interested in calculating the trace and determinant of $\mathbf{H}_{\tilde{c}}$ at $D_1 = D_2 = D^*$, with $\nabla \tilde{c}|_{D_1 = D_2 = D^*} = 0 = c'(D^*)$. For the sake of brevity in notation, we will omit dependence on $D^*$ in the following equations of this subsection. All symbols to appear are assumed to be calculated at $D^*$ (e.g. $D^* \to D$, $r''(D^*) \to r''$ and so on).

Note that $c' = 0$ implies $r'D = -r$ and also $c'' = \frac{8r}{c^3}(D^2 r'' - 2r)$. With this, $\partial^2_{D_1} \tilde{c} = \partial^2_{D2} \tilde{c} = \frac{4r}{c^3}(D^2 r'' - r)$, hence

$$\text{tr}(\mathbf{H}_{\tilde{c}}) = \frac{8r}{c^3}(D^2 r'' - r). \tag{36}$$

Whereas $\partial^2_{D_1, D_2} \tilde{c} = \partial^2_{D_2, D_1} \tilde{c} = -\frac{4r^2}{c^3}$ leads to

$$\det(\mathbf{H}_{\tilde{c}}) = \frac{16r^2}{c^6}(D^2 r'' - r)^2 - \frac{16r^4}{c^6} \tag{37}$$

$$= \frac{2rD^2}{c^3} r'' c''. \tag{38}$$

In the main text we only focus on the sign of the trace and determinant, which depend on $D^2r'' - r$ and $r''c''$, respectively.

### 5 Numerical Exploration

We let the resident population establish a traveling wave profile of speed $c_1 = 2\sqrt{r_1 D_1}$. After sufficient time ($t = t_{inv} = 20$), we introduce a mutant with speed $c_2 = pc_1$, with $p \in \mathbb{R}$. The initial mutant density is centered at a given point in space, $x^*$, where the density of the resident population is given by $u_1(x^*, t_{inv})$. Then, we set $u_2(x, t_{inv}) = u_1(x, t_{inv})$ for $x \in [x^* - 1/2, x^* + 1/2]$ and $u_2(x, t_{inv} = 0)$ otherwise, and set $u_1(x, t_{inv}) = 0$ for $x \in [x^* - 1/2, x^* + 1/2]$. From there, we let the system evolve in time, and access wether mutant population densities are eventually much higher than resident populations at the leading edge, if so, we state that mutants invade successfully, and say they fail otherwise (we set a maximum number of time-steps for the simulation). We perform that analysis for different values of $p = c_2/c_1$ and $u_1(x^*, t_{inv})$, leading to figure 2.

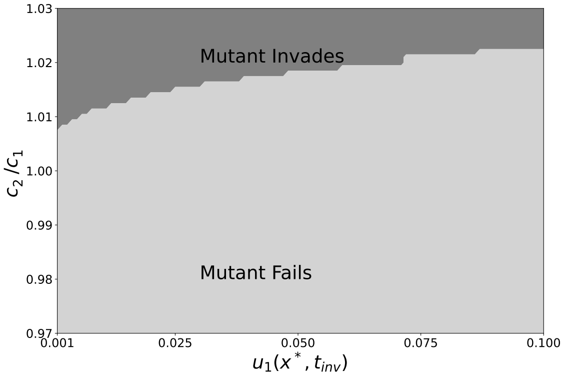


Figure 2: Numerical Invasion analysis. We test different values of mutant population speed at different resident population density levels. The result indicates the $c_2 > c_1$ is a reasonable criteria for mutant invasion
